# Tempo and mode of gene expression evolution in the brain across Primates

**DOI:** 10.1101/2021.04.21.440670

**Authors:** Amy L. Bauernfeind, Trisha M. Zintel, Jason Pizzollo, John J. Ely, Mary Ann Raghanti, William D. Hopkins, Patrick R. Hof, Chet C. Sherwood, Courtney C. Babbitt

**Affiliations:** Department of Neuroscience, Washington University School of Medicine, St. Louis, MO 63110, USA; Department of Anthropology, Washington University in St. Louis, St. Louis, MO 63130, USA; Department of Biology, University of Massachusetts Amherst, Amherst, MA 01003, USA; Molecular and Cellular Biology Graduate Program, University of Massachusetts Amherst, Amherst, MA 01003, USA; Alamogordo Primate Facility, Holloman Air Force Base, NM 88330, USA; Department of Anthropology, School of Biomedical Sciences, and Brain Health Research Institute, Kent State University, Kent, OH 44242, USA; Department of Comparative Medicine, Michale E. Keeling Center for Comparative Medicine, The University of Texas M D Anderson Cancer Center, Bastrop, TX 78602, USA; Nash Family Department of Neuroscience and Friedman Brain Institute, Icahn School of Medicine at Mount Sinai, New York, NY 10029, USA; New York Consortium in Evolutionary Primatology, New York, NY 10124, USA; Department of Anthropology and Center for the Advanced Study of Human Paleobiology, The George Washington University, Washington, DC 20052, USA

## Abstract

Primate evolution has led to a remarkable diversity of behavioral specializations and pronounced brain size variation among species ^1,2^. Gene expression provides a promising opportunity for studying the molecular basis of brain evolution, but it has been explored in very few primate species to date e.g. ^3,4^. To understand the landscape of gene expression evolution across the primate lineage, we generated and analyzed RNA-Seq data from four brain regions in an unprecedented eighteen species. Here we show a remarkable level of variation in gene expression among hominid species, including humans and chimpanzees, despite their relatively recent divergence time from other primates. We found that individual genes display a wide range of expression dynamics across evolutionary time reflective of the diverse selection pressures acting on genes within primate brain tissue. Using our sample that represents an unprecedented 190-fold difference in primate brain size, we identified genes with variation in expression most correlated with brain size and found several with signals of positive selection in their regulatory regions. Our study extensively broadens the context of what is known about the molecular evolution of the brain across primates and identifies novel candidate genes for study of genetic regulation of brain development and evolution.

---

Primates are distinguished from other mammals by their large brains relative to body size ^5^. Among the diversity of primate species, there is remarkable variation in behavioral specializations, including differences in social structure, spatial, dietary and visual ecology, and locomotion ^2^. Despite the impressive array of cognitive attributes displayed by primates, studies investigating various aspects of brain evolution tend to sample from a small number of species to address questions of how humans are unique. In large part, the emphasis on human brain evolution is warranted. Humans are unmatched in possessing exceptionally large brains and unparalleled cognitive abilities, such as language ^6,7^. While valuable, the limited number of species included in prior research does not provide a comprehensive perspective of the context in which the human brain evolved within the diversity of primates.

Although researchers have used a variety of approaches to assess whether the human brain is unique ^8^ and how it might have evolved ^9,10^, the potential for using gene expression studies to evaluate patterns of brain evolution in primates has not yet been met. Upon observing the remarkable similarity between human and chimpanzee protein sequences, King and Wilson ^11^ proposed that the basis of the physical and behavioral phenotypic differences between these two species must be found in changes within gene regulatory regions that drive expression. Previous studies have explored how changes in regulatory regions can influence gene expression but have often sampled various organs from species across broad spans of evolutionary time, such as mammals or vertebrates ^12,13^. In studies focusing on gene expression in primate brain tissues, research has mostly focused on the neocortex and cerebellum in human, chimpanzee, and rhesus macaque ^3,4,14-16^. However, new insights can be gained by sampling at greater neuroanatomical resolution from a broader array of primates. Examining gene expression of the brain from a more comprehensive landscape empowers novel inquiry in primate brain evolution, including questions pertaining to the sources of variation that drive expression differences, rates of expression change across the primate phylogenetic tree, and genes that correlate with brain size across primates.

In the current study, we sampled prefrontal cortex (PFC), primary visual cortex (V1), hippocampus (HIP), and lateral cerebellum (CBL) from 18 primate species, the broadest diversity of primates sampled in any study of gene expression in the brain to date, including species from several rarely-studied lineages. Our dataset represents 70-90 million years ^17^ of primate evolution, providing a more thorough understanding of the evolution of gene expression across primates and allowing for an unprecedented view of how gene expression in the brain has changed over time across all major clades of primate phylogeny.

## Results

### Variation in brain gene expression across the primate phylogeny

To understand how gene expression in the brain has evolved across the primate lineage, we generated and analyzed RNA-seq data from 18 primate species (including 5 hominoids, 4 cercopithecoids, 4 platyrrhines, and 5 strepsirrhines) across 4 brain regions, including PFC, V1, HIP, and CBL (**Fig. 1, Supplementary Table 1**). Not all of the sampled species have publicly available genomes, so transcriptomes and gene models were assembled *de novo* ^18^ (see **Supplementary Information**). We quantified expression of 15,017 orthologs within hominoids, and 3,432 one-to-one orthologs across all 18 species (**Supplementary Table 2**).

**Fig. 1.**
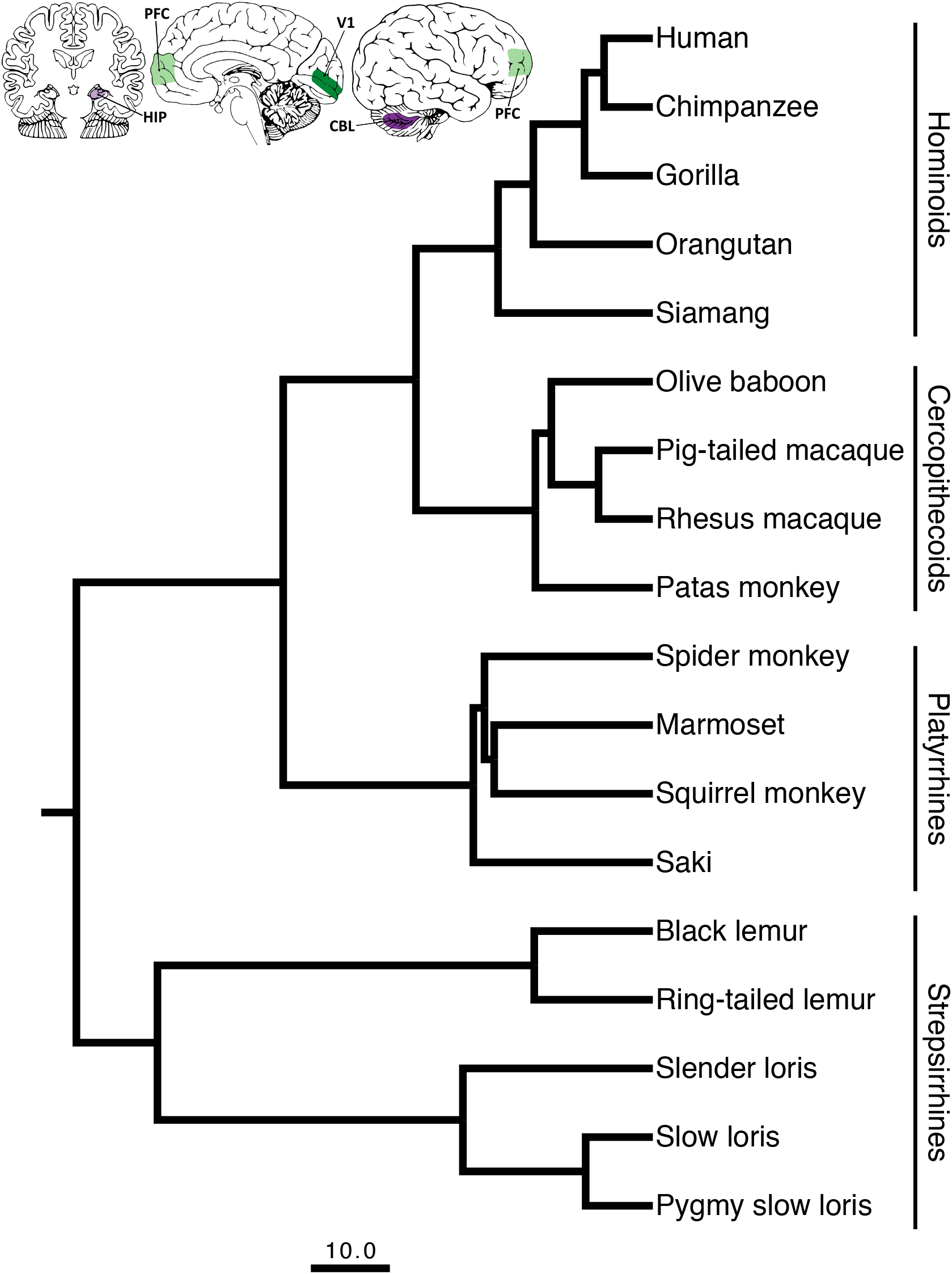
Primate phylogeny showing the eighteen species sampled in this study. The scale bar for the branch lengths represents 10 million years of evolution. The phylogenetic tree is a consensus tree of 1000 iterations produced from *10kTrees* v.3 (https://10ktrees.nunnlab.org) based on data from GenBank. The insets demonstrate the approximate locations of the four brain regions sampled on a coronal section, midsagittal view, and lateral view (displayed left to right, respectively) of a schematized adult human brain.

Variation in interspecific mammalian gene expression has been shown to be less pronounced than that observed across samples from different organs, reflecting the diversity of underlying organ physiology ^12,19^. Furthermore, it has been reported that the rate of gene expression divergence evolved more slowly within the cerebral cortex and cerebellum compared to other organ systems from developmentally distinct germ layers^4,12^. To explore the variability of gene expression from distinct regions of the brain across our broad sampling of primates, we constructed a pairwise distance matrix of the 500 most variable protein-coding genes based on standard deviation of expression across samples (**Methods**). This subset of genes was enriched with glycoproteins, signal peptides, and plasma membrane proteins, with roles in immune function, molecular trafficking, and cell signaling. Using this distance matrix, we performed a principal coordinates analyses (PCoA) on data from all brain regions. Because our samples represent disparate regions of the same organ, we expected less variation to be attributed to brain region than primate species or taxa, reflecting the similarity in physiology of brain tissues. Unsurprisingly, the variation from our complex gene expression dataset is represented across multiple axes of the PCoA (**Supplementary Table 3**).

We plotted the first three axes and created polygons around data derived from common taxa (**Fig. 2a-c**) or region (**Fig. 2d-f**). As predicted, taxon assignment explains a large amount of variation to the dataset, with clear trends emerging independent of brain region. We find the greatest divergence in expression patterns among hominoid and strepsirrhine species, while there is more similarity observed among cercopithecoids and platyrrhines. The hominoids, displaying the greatest level of diversity of any primate phylogenetic group, demonstrate variation that is particularly apparent along Axis 1 and largely driven by human and chimpanzee expression patterns (**Extended Data Fig. 1**). Strepsirrhines also exhibit a large amount of variation, especially apparent along Axis 2, which can largely be attributed to the three species of lorises. When Axis 1 and 2 of the PCoA are plotted on the same bivariate plot (**Fig. 2a**), the hominoids display more variation than the strepsirrhines by about 24% (**Supplementary Table 4)**. However, a large portion of the variation in the strepsirrhines is attributed to evolutionary divergence over about 63 million years (since the last common ancestor of lemurs and lorises), whereas the variation within hominoids has largely accrued over only 9 million years (since humans and chimpanzees shared an ancestor with gorillas). The hominoid and strepsirrhine samples represent similar variation in terms of sex and life stage, suggesting that these factors do not account for the variability seen in these taxa. Therefore, a remarkable finding of this analysis is how much variation is represented by hominoids, despite the fact that this lineage represents a much shorter evolutionary divergence time.

**Fig. 2.**
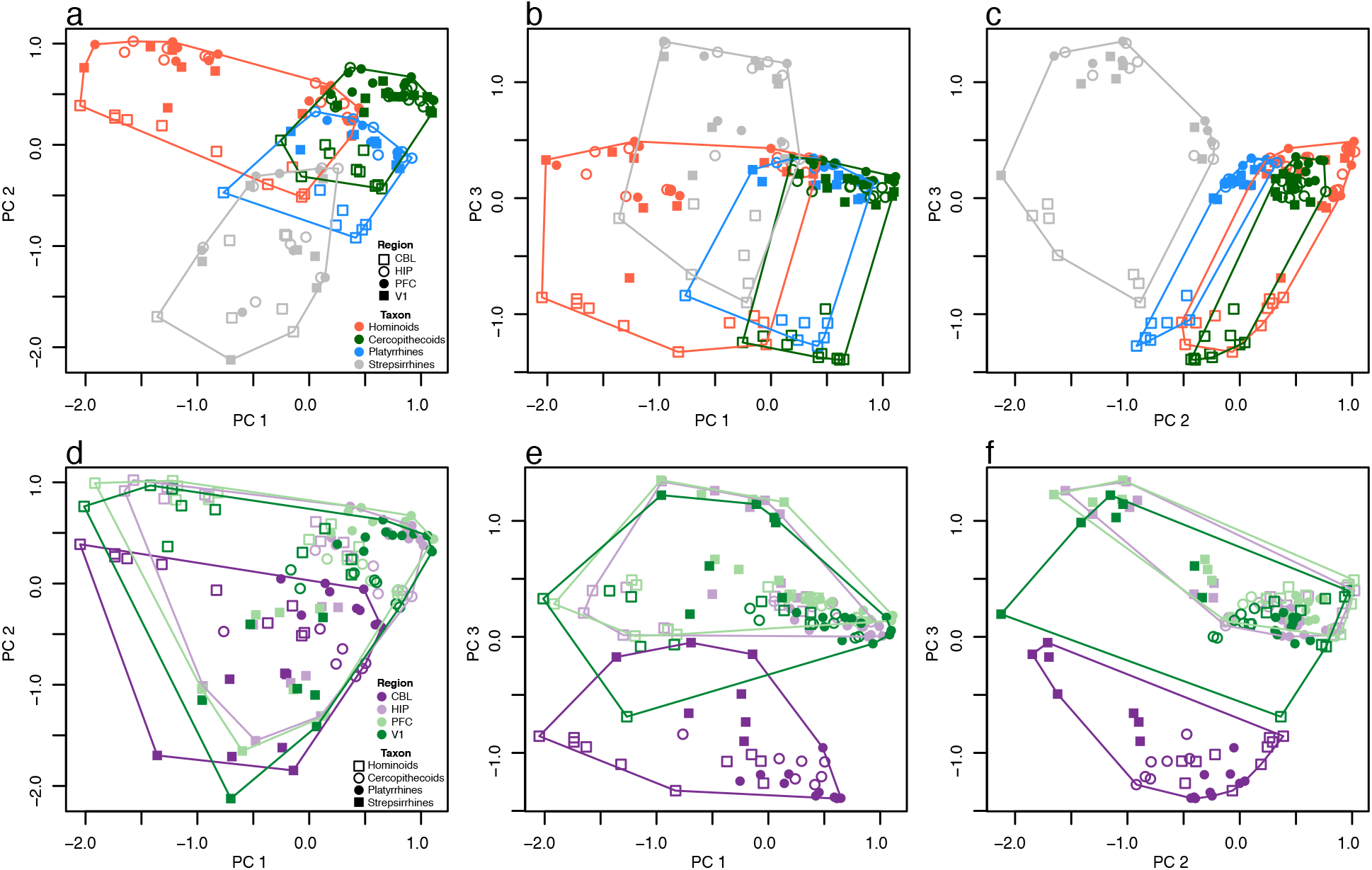
Patterns of brain gene expression across primates. The first three axes of a principal coordinates analysis (PCoA) are plotted in both rows but have different symbols and colors to emphasize expression patterns specific to taxa (upper row, a-c) and regions (lower row, d-f). Polygons in each plot surround the data points for taxa (upper row) and regions (lower row). Axes 1, 2, and 3 represent 12.8, 10.3, and 9.4% of variance, respectively.

We observed trends in gene expression by brain region that are predominantly seen along Axis 3 (**Fig. 2e-f)**. Here, we observe that the variation attributed to CBL is distinguished from that of PFC, V1, and HIP, which are very similar in their distributions. The fact that CBL differs in its pattern of gene expression from other brain regions ^15,20,21^ is not surprising given that it is the only sampled region that develops from a different part of the embryonic neural tube (namely, the hindbrain vs. forebrain) and exhibits a neuronal packing density (predominantly glutamatergic granule cells) that far exceeds these other brain regions^22-24^). Enrichments of cerebellar tissue reveal expression changes to semaphorin genes between humans and chimpanzees, and between humans and most other primate species with deeper evolutionary relationships (**Supplementary Table 5**). Notably, semaphorin genes have been found to be enriched in human-specific enhancer regions ^25^ and have been shown to mediate the cell-cell interactions necessary for axon guidance ^26^. Although the expression levels of semaphorin genes contribute to the distinction between human CBL compared to other species, these genes are not enriched in contrasts of brain regions other than the CBL indicative of region-specific differential expression.

To determine whether gene expression profiles can reconstruct known phylogenetic relationships among primates, we built expression phenograms (**Methods**) for all brain regions combined (**Extended Data Fig. 2**) and each region separately (**Extended Data Fig. 3**). Samples that were derived from individuals of the same species tended to be grouped together, regardless of brain region, revealing that inter-individual differences are minor compared to other sources of variation. Gene expression profiles also replicated the phylogenetic relationships of closely related species (e.g. humans and chimpanzees; pig-tailed and rhesus macaques) when all regions were considered, but these relationships became less phylogenetically structured in the phenograms constructed using expression data from individual brain regions. All phenograms accurately represent cercopithecoids and strepsirrhines as monophyletic groups; however, expression data produces paraphyletic groups of hominoid and platyrrhine species. This result potentially reflects the fact that taxa with longer periods of independent evolution (i.e., strepsirrhines) are more likely to show divergent patterns of gene expression than more closely related groups. Meanwhile, more dense sampling of cercopithecoids (3 individuals per species), permits a fairly accurate reconstruction of this taxon.

Across all one-to-one orthologs represented in the sampled primates, we found that ∼15-20% of genes show differential expression across species (q-value < 0.05). As expected, the relative amount of differential expression increases over evolutionary time, both between species and clades (**Fig. 3**, Human-Chimpanzee through Human-Lemur columns). However, both the species and clade-wise comparisons show larger numbers of differentially expressed (DE) genes in the comparisons when strepsirrhines are included in the contrast, a result of the more than 60 million years of independent evolution of this taxon. By region, there are similar numbers of DE genes in CBL, HIP, and PFC, with far fewer changes in gene expression seen in the V1 across all contrasts. Such a finding may be expected, given the high ratio of genes in V1 that have been found to display conserved expression, even between human and mouse samples ^27^.

**Fig. 3.**
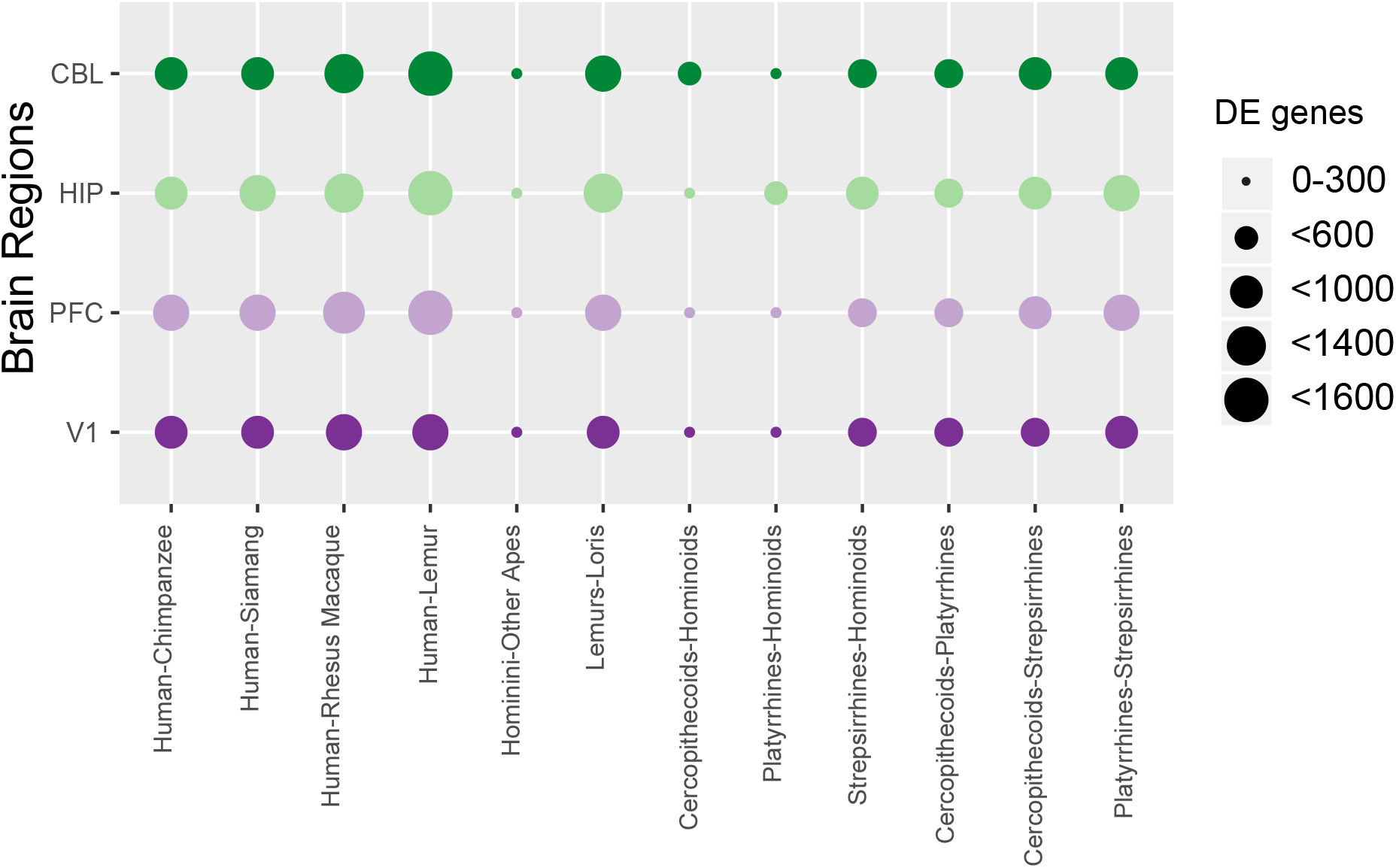
Gene counts of DE genes between species and clades. Each row represents one of the four brain regions examined. The size of the circle represents the number of DE genes seen at q < 0.05 (5% FDR). The comparisons on the left are between exemplar species or sets of species, comparisons on the right are between clades of primates.

When comparing gene expression in PFC of humans relative to other primates, humans PFC shows an enrichment of metabolic processes, including “regulation of cellular metabolic process” and “regulation of macromolecule metabolic process” (**Supplementary Table 5**). Comparing human and chimpanzee PFC reveals that categories that support neural growth and development (e.g., “neuron projection morphogenesis”, “cell morphogenesis involved in neuron differentiation”), gene regulation, and metabolic processes are enriched in human PFC relative to chimpanzees (**Supplementary Table 5)**. The enrichment of neuronal projection and connective processes may be related to the enhanced white matter connectivity of human brains as compared to those of chimpanzees ^28-31^.

Our PCoA analyses showed that gene expression in brain regions sampled from the lorises (i.e. the slender loris, slow loris, and pygmy slow loris) diverged from other strepsirrhines, and other primates more generally (**Extended Data Fig. 1**). When strepsirrhines are compared to other primates in differential expression analyses, transcription factors and other genes involved in gene transcription and translation and multiple biosynthetic pathways involved in cellular metabolism are among the categories of DE genes (**Supplementary Table 5)**.

### Evolutionary rates of expression change across clades and brain regions

Previous studies have used a variety of different approaches to model gene expression changes over time ^12,32^. Here, we used a recently described model to analyze neutral and conserved processes as determined by changing gene expression levels ^33^. We found that individual genes exhibit wide variation in expression dynamics across the primate lineage (**Fig. 4a)**. Enrichments for genes showing low variation or stabilizing selection (q = 0.05) reveal categories related to transport and cellular localization (GO Biological Processes, **Supplementary Table 6**). In contrast, genes that are less constrained or neutrally evolving (q > 0.05) have a number of processes related to neuron morphogenesis, plasticity, and cell death. Chen et al.^33^ observed that, unlike sequence evolution, gene expression is not linear across evolutionary time but a saturation point in pairwise comparisons of gene expression is reached due to stabilizing selection pressures. Here, we find that pairwise expression differences between humans and the other species increasingly diverge with evolutionary distance in all brain regions sampled (**Fig. 4b**); however, these pairwise comparisons do not seem to saturate with evolutionary time across the primate comparisons ^33^. We note that the saturation of pairwise expression differences from humans may be found at a phylogenetic node ancestral to primates (**Extended Data Fig. 4**).

**Fig. 4.**
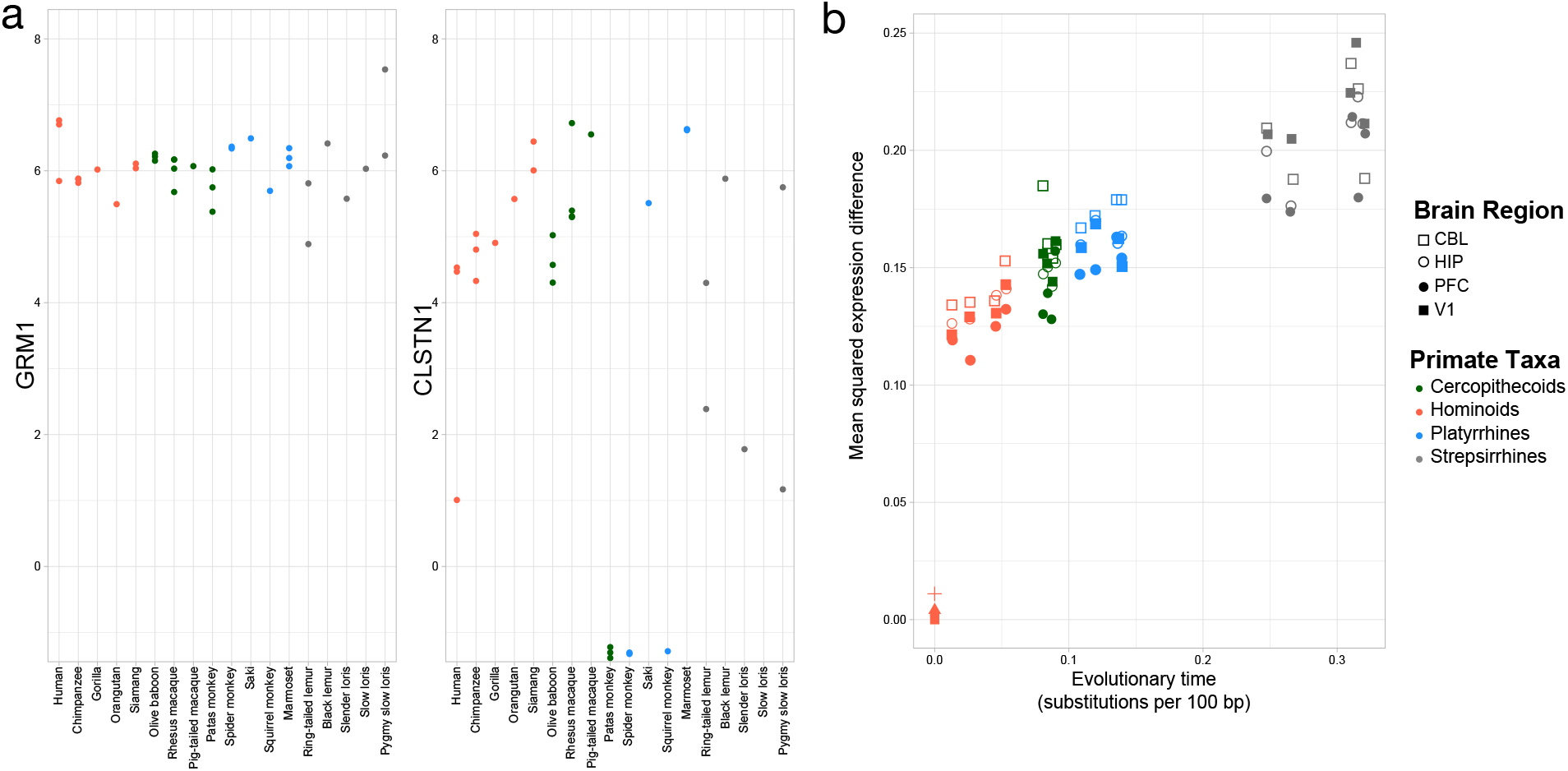
Rates of change over genes and evolutionary time. a. Exemplar genes that show constraint (left panel) and variation (right panel) across primates (colors as in Figure 2). b. Mean squared expression difference plotted by evolutionary distance across all orthologs that were expressed. Shapes denote the four brain regions, and the colors represent the four major primate clades represented in our samples.

### Correlation of gene expression to brain size and their change over evolutionary time

Because an increase in absolute brain size is one of the most striking characteristics of humans, we asked what subset of genes is correlated with this important biological trait across the primate tree. Our dataset provides a unique opportunity to evaluate this question, since average brain size across our study species varies by ∼190-fold (**Supplementary Table 7**). Using multivariate analysis (Methods), we defined gene lists most strongly correlated with brain size in each brain region (**Supplementary Table 8**). Results indicate that the same genes rank among those with the strongest positive correlation in PFC, V1, and HIP. CBL also shares some of these same genes but includes more variation among the genes most strongly correlated to overall brain size than the other three brain regions (**Fig. 5, Supplementary Table 8**), potentially reflecting the more recent expansion of the CBL relative to the rest of the neocortex ^34^. To explore possible adaptive changes in regulation of the genes that correlated with brain size, we analyzed the putative regulatory regions for signatures of positive selection. We tested for an excess of change in 5’ proximal gene regulatory regions^35^ as compared to neutral proxy regions, which is suggestive of positive selection. It is important to note that this test is for ancient, not modern, signatures of positive selection since individual mutations have had time to be accumulate in regulatory regions and introns. Of 207 genes with correlations with brain size tested, we found eight with signatures of positive selection (p < 0.05) (**Table 1, Supplementary Table 9, 10**). *Pbx1*, is a member of the homeobox family of transcription factors, shown to promote regional and laminar patterning of the cerebral cortex ^36^. In postnatal rat brains, it is expressed within the subventricular zone, olfactory bulb, and along the migratory path between these two areas, suggesting a role in neurogenesis and the migration of neurons ^37^. *Pax2* also participates in the patterning of the cerebral cortex and is expressed in a decreasing rostral to caudal gradient in mice from the earliest stages of development ^38,39^. Interestingly, the amino acid sequence of *Pax2* itself is 87% the same in mice compared to zebrafish, consistent with an evolutionarily conserved role in the development of the rostral aspect of the cerebral cortex^39^. *Pten* is a key regulator of signal transduction, including acting as an antagonist of the PI3K/AKT pathway which controls cell survival and dendritic arborization^40^. As such, *Pten* knockout and hybridized mice develop improper numbers of neurons, neuronal size, and extent of dendritic branching ^41^. The list of genes correlated with brain size provides novel candidates for exploration of genes on primate brain growth and development.

**Figure 5.**
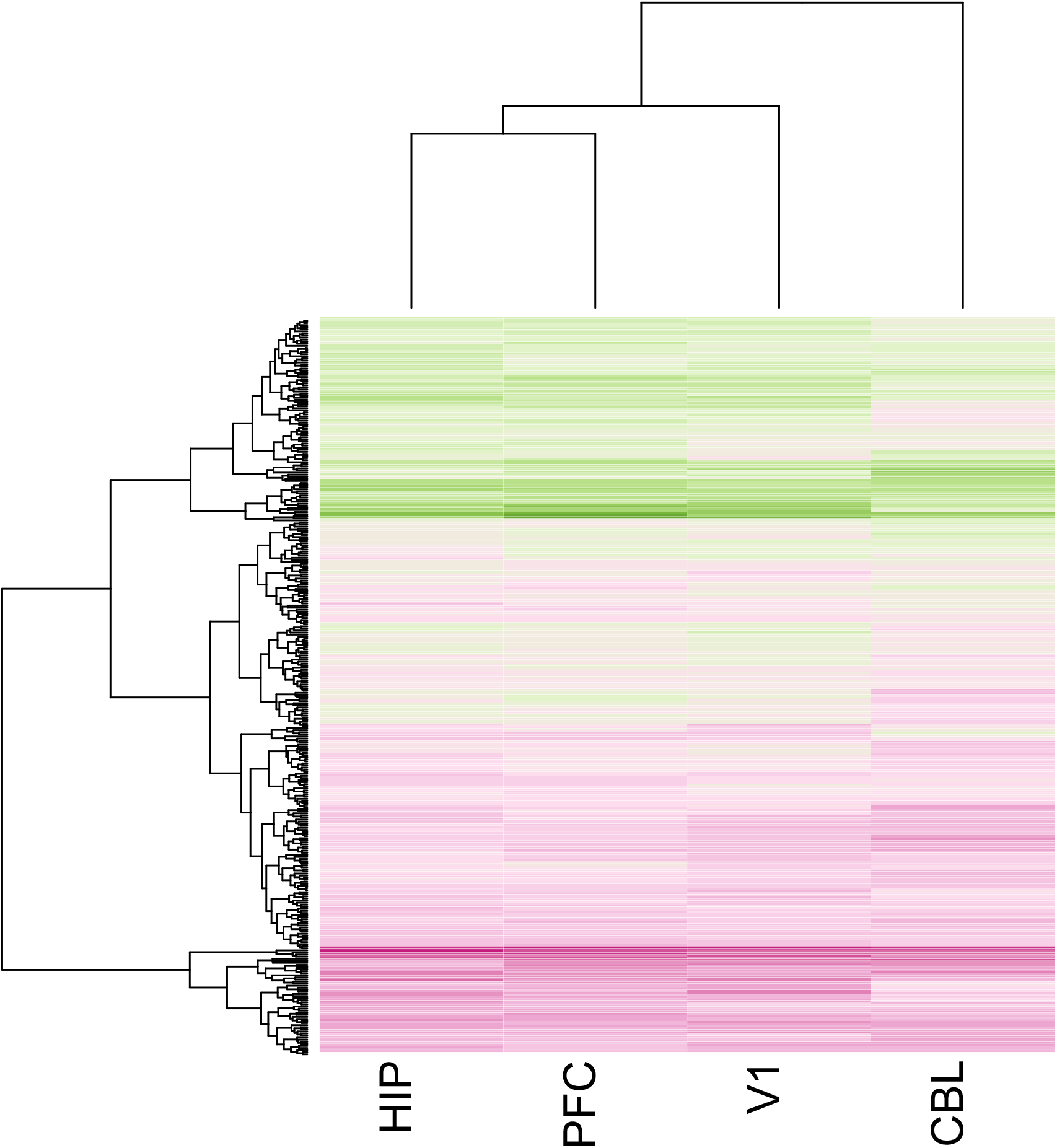
Gene correlated with brain size by region. A clustering and heatmap of the residuals from PC2 of genes for the four regions examined (V1, HIP, PFC, and CBL).

## Discussion

We performed RNA-seq on 4 brain regions from 18 primate species, representing the broadest sampling for any gene expression study in primate brain tissue to date. Through more representative sampling of primate species, we found substantial variation in gene expression levels within the hominoid and strepsirrhine lineages, with the hominoid diversity particularly impressive due to the recent divergence of this taxon. The considerable diversity of gene expression in human and chimpanzee brain tissue has profound implications for understanding the distinct evolutionary processes that have acted upon the brains of the ancestral species of these two lineages. We observed a wide variety of expression dynamics of individual genes in the pairwise comparisons of humans to other primate species. However, less constrained, neutrally evolving patterns appeared to be the most prevalent pattern in each brain region studied, preventing a saturation point of stabilizing pressures to be reached within Primates with increased phylogenetic distance from humans. Lastly, we identified genes that are correlated with brain size across all major primate taxa, providing candidates for further inquiry.

Our deeper analysis of gene expression has revealed evolutionary patterns that were inaccessible with more limited sampling of primate brain tissue. We anticipate that the candidate genes and data provided by this study will serve as a resource for many other lines of inquiry into human and non-human primate brain evolution.

## Methods

The sample includes brain tissue from human and nonhuman primates. All samples were obtained from adult individuals free from known neurological disease. If available, the right hemisphere was preferentially sampled. Human brain samples were obtained from the National Institute for Child Health and Human Development Brain and Tissue Bank for Developmental Disorders at the University of Maryland (Baltimore, MD). Chimpanzee brain tissue was obtained from the National Chimpanzee Brain Resource (https://www.chimpanzeebrain.org, supported by NIH grant NS092988). All other sources of brain tissue are listed in **Supplementary Table 1** and other details about sampling can be found in the **Supplementary Information**.

We extracted high-quality RNA and prepared poly-A pulldown libraries using standard protocols. Illumina libraries were sequenced on a NextSeq500 and transcriptomes were assembled using the Trinity package ^42^. Blast was used to assign orthologs across species, and counts were normalized using RSEM ^43^ and edgeR ^44^.

We performed a principle coordinates analyses (PCoA) based on a pairwise distance matrix of all 137 samples. The distance matrix was comprised of the top 500 most variably expressed protein-coding genes by standard deviation across samples. Pairwise distances were calculated by log2 fold change, providing a symmetrical representation of the expression ratio centered around 0 (ie. log2(2) = 1 while log2(0.5) = −1). Although variation is represented across more than 20 axes (**Supplementary Table 3**), the first three axes were plotted to compare patterns across primate taxa and brain region sampled (**Figure 2**). **Extended Data Figure 3** is the same graph except using an array of colors that allow the data from each individual species to be visualized.

Putative signatures of positive selection in promoters were inferred by comparing substitution rates in promoters to rates in nearby neutrally evolving portions of the genome. The test for selection was performed using modified code from Haygood et al. 2007 ^35^, and run using HyPhy software ^45^. Data availability: Sequencing data have been deposited in the Short Read Archive: BioProject PRJNA639850; the link for reviewers is: https://dataview.ncbi.nlm.nih.gov/object/PRJNA639850?reviewer=136ah9q3o5ok7a7gq8jd4hhdfc

## Acknowledgements

We would also like acknowledge our funding from NSF BCS-1750377, Wenner Gren Foundation, James S. McDonnell Foundation (220020293), NSF INSPIRE (SMA-1542848), and NIH (NS-092988).

## Author Contributions

CCB, JJE, MAR, WDH, PRH, CCS, and ALB designed the study. CCB, TMZ, JP, and ALB analyzed the data. CCB and ALB wrote the manuscript.

## Author Information

The authors declare no competing interests. Correspondence and requests for materials should be addressed to.

## Tables

**Table 1**. Top genes from the brain size analysis that show signatures of positive selection in their proximal 5’ regulatory regions. Selection tests were performed on sequences with available overlapping human-chimpanzee-macaque alignments. Tests were performed excluding overlapping coding regions (promoter non-coding only) that may have regions with constrained nucleotide substitution, and also on sequences that include these overlapping coding regions (promoter full sequence). This method compares rates of substitution in promoters to surrounding neutrally evolving regions and uses a likelihood ratio test to give a p-value for agreement of substitution rate in tested regions compared to a null model representing no or negative selection. The phyloP test models nucleotide changes across branches of a phylogeny and uses prior and posterior distributions to generate p-values for selection on a branch of the phylogeny. The number of nucleotides under selection was tested with the phyloP LRT method that computes base-by-base p-values for accelerated substitution in promoters.

## EXTENDED DATA

**Extended Data Figure 1.**
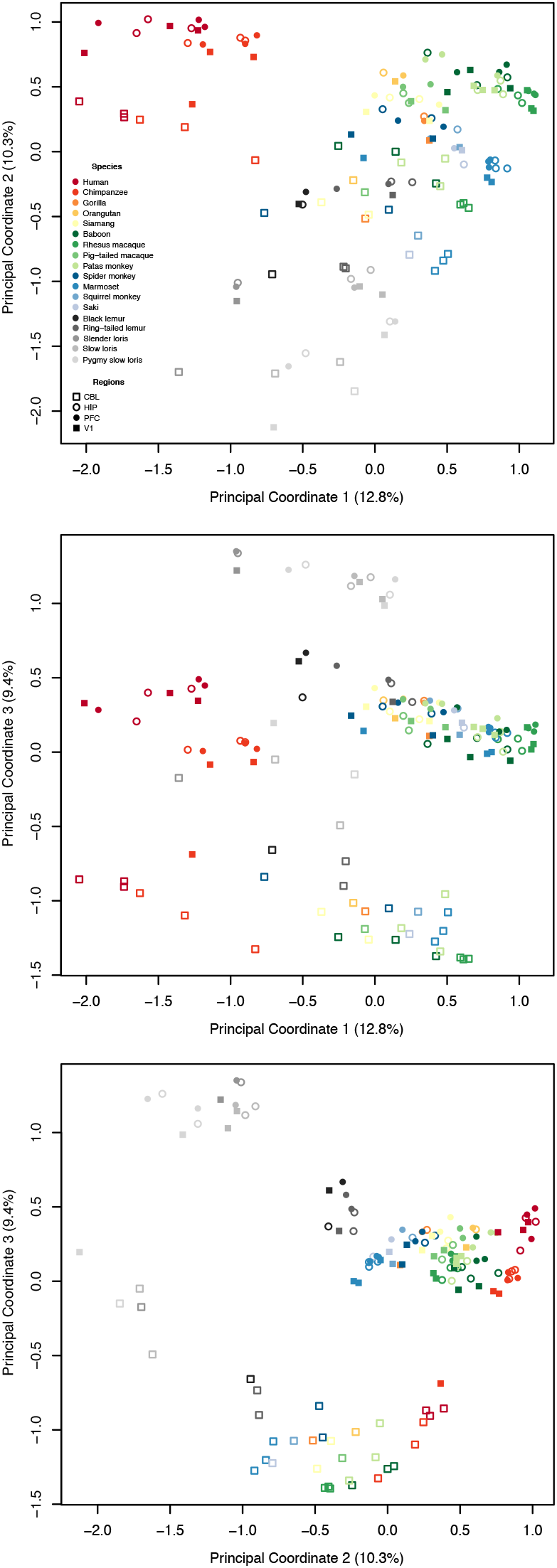
The first three axes of the PCoA are plotted in three bivariate plots. This is the same plot as Figure 2 of the main text except here each species is plotted as a different color.

**Extended Data Figure 2.**
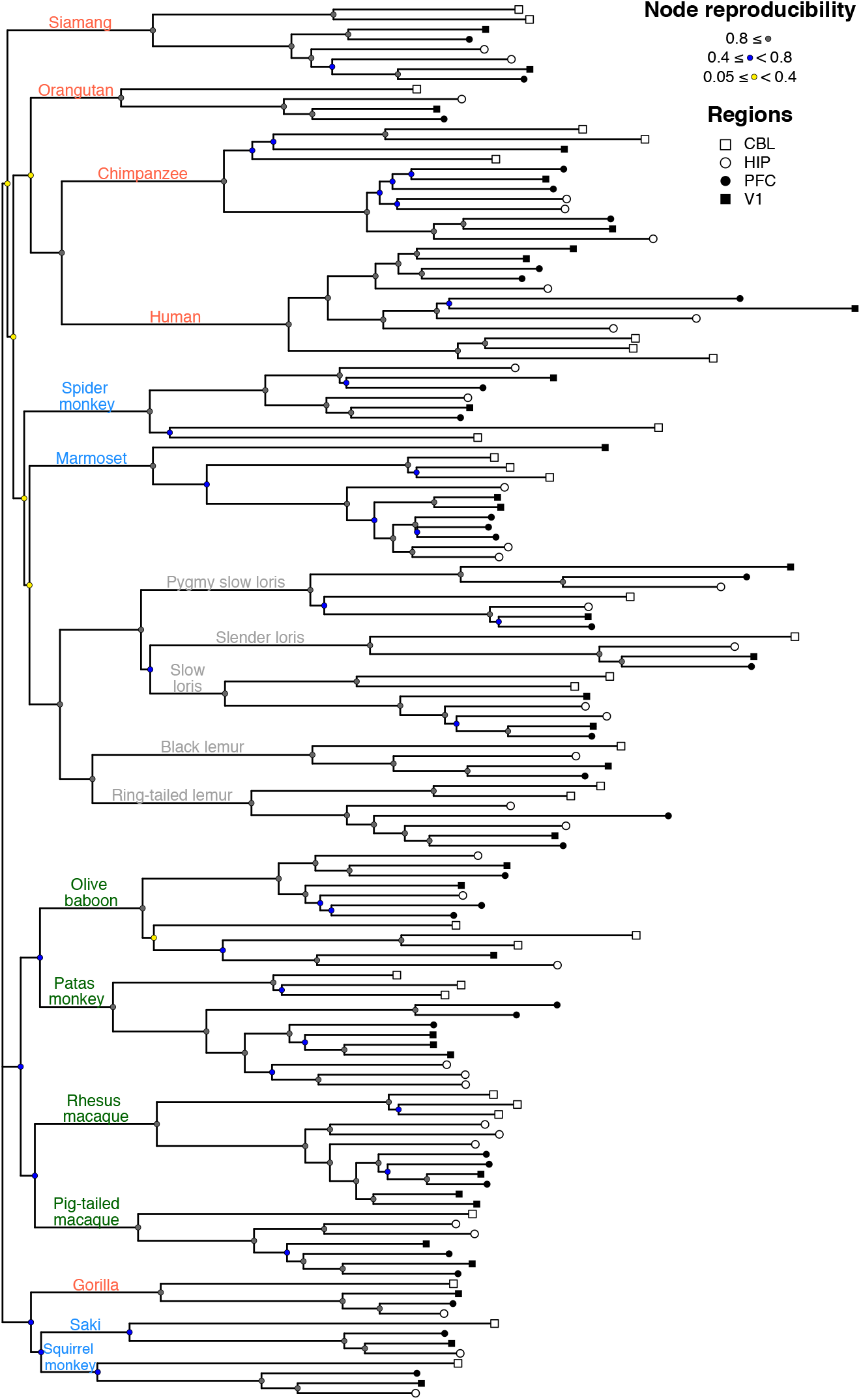
Gene expression phenogram of all sampled data. The phenogram was constructed from neighboring-joining tree estimation based on log2 fold-change distances of the 500 most variable genes based on standard deviation (the same distance matrix that constructed the PCoA). Reproducibility of the nodes of the tree were estimated using a bootstrap analysis of 1000 iterations. Species common names are color coded to indicate taxa (hominoid, pink; cercopithecoid, green; platyrrhine, blue; strepsirrhine, gray).

**Extended Data Figure 3.**
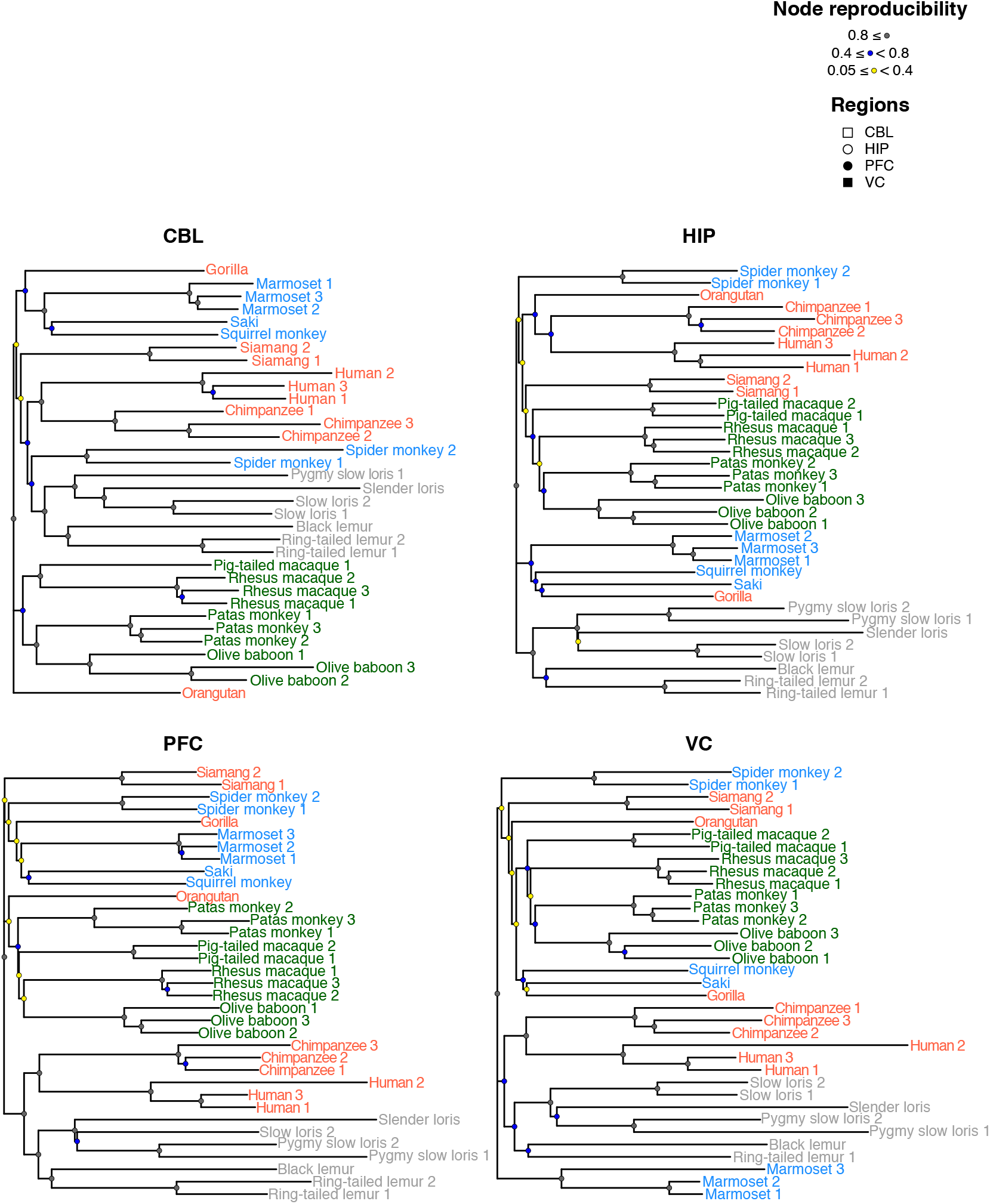
Gene expression phenograms by region. The phenograms were constructed using the same techniques as Extended Data Figure 2. Species common names are color coded to indicate taxa (hominoid, pink; cercopithecoid, green; platyrrhine, blue; strepsirrhine, gray).

**Extended Data Figure 4.**
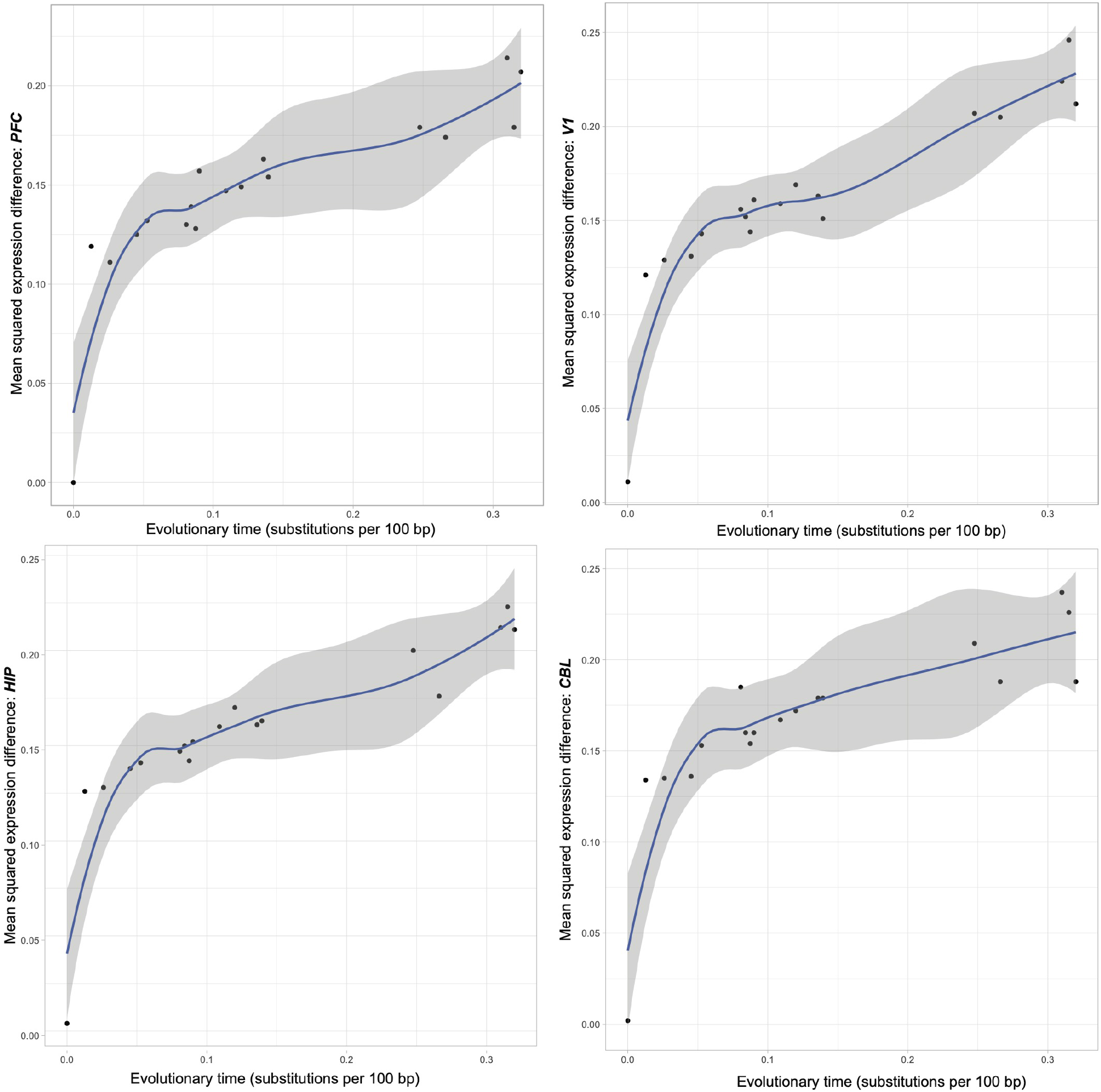
Region-specific plots of mean square expression differences over evolutionary time for each of the four brain regions analyzed.

**Extended Data Figure 5.**
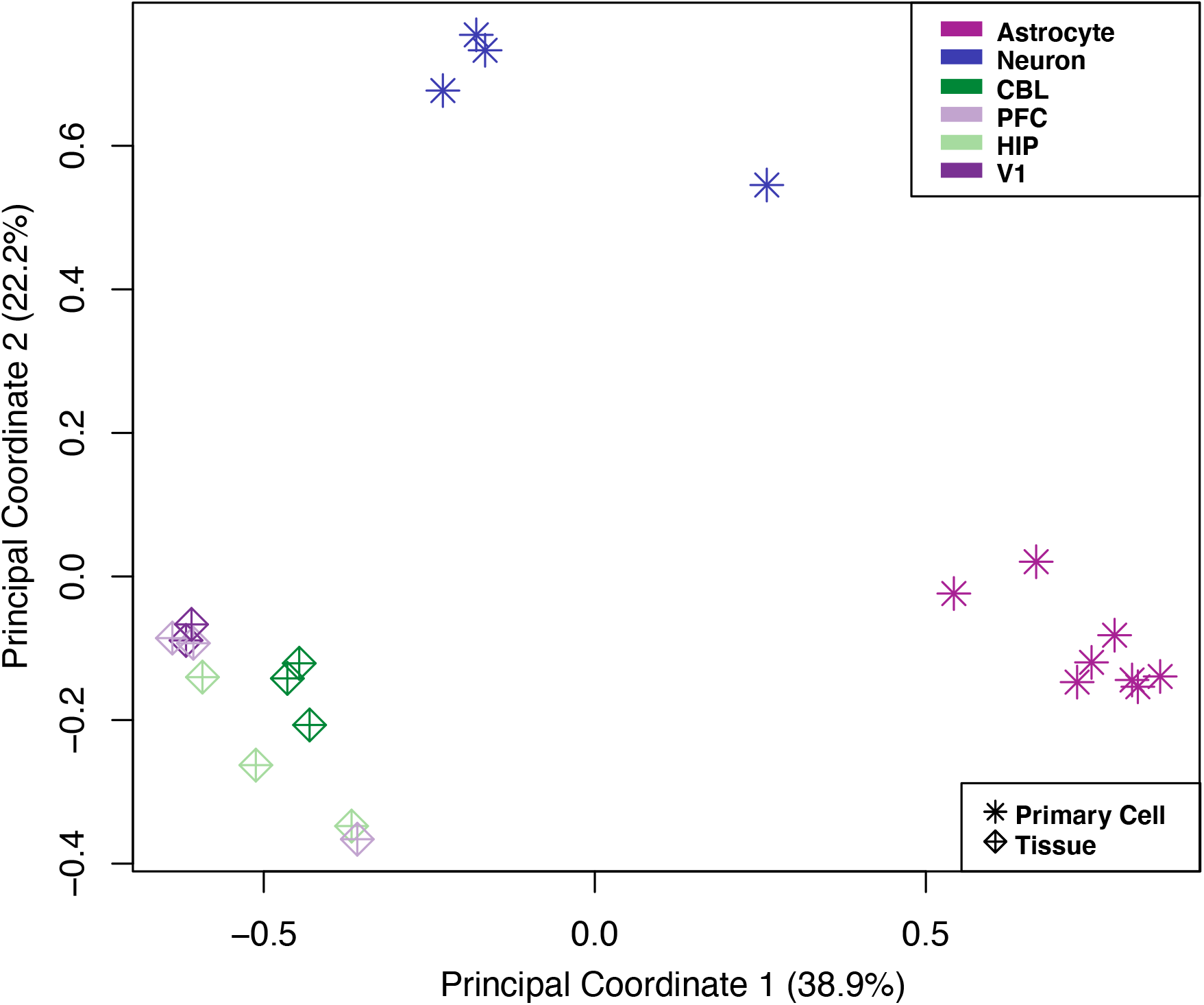
PCoA of expression data from human samples of all four brain regions and primary neurons and astrocytes. Symbols delineate the current study’s tissue samples from primary cell data. Colors distinguish cell type or brain region. Tissue level human samples from all regions do not exhibit biased expression for either neurons or astrocytes.

## SUPPLEMENTARY INFORMATION

### Biological sample collection and RNA extraction

All nonhuman primates had been cared for according to Federal and Institutional Animal Care and Use guidelines. All nonhuman primates died of natural causes or were euthanized for humane reasons to reduce suffering from terminal illness. All individuals sampled were free from known neurological conditions. Human and nonhuman brains were frozen after a postmortem interval of no more than 8 h and stored at −80°C.

From each individual, we sampled four regions of the brain, including prefrontal cortex (PFC), primary visual cortex (V1), hippocampus (HIP), and cerebellum (CBL). PFC was sampled from the frontal pole, corresponding to Brodmann’s area 10 in humans. In other primates, the PFC region sampled more broadly encompassed prefrontal cortical areas but was limited to the most anteriorly projecting part of the frontal pole. All V1 samples were dissected around the calcarine sulcus to include primary visual cortex (Brodmann’s area 17). Samples from both PFC and V1 contained all cortical layers and a small amount of underlying white matter (<10%). The HIP was sampled from the medial aspect of the temporal lobe and included all hippocampal subfields. The CBL was sampled from the most laterally projecting region of lateral hemisphere in all primates. In humans, the CBL region corresponded to Crus I or Crus II. CBL samples contained all layers of cerebellar cortex and a small amount of underlying white matter (<10%). Each sample was briefly homogenized using a Tissuelyzer (Qiagen), and the total RNA isolated using RNAeasy kit (Qiagen) with a DnaseI treatment.

**Table S1.**
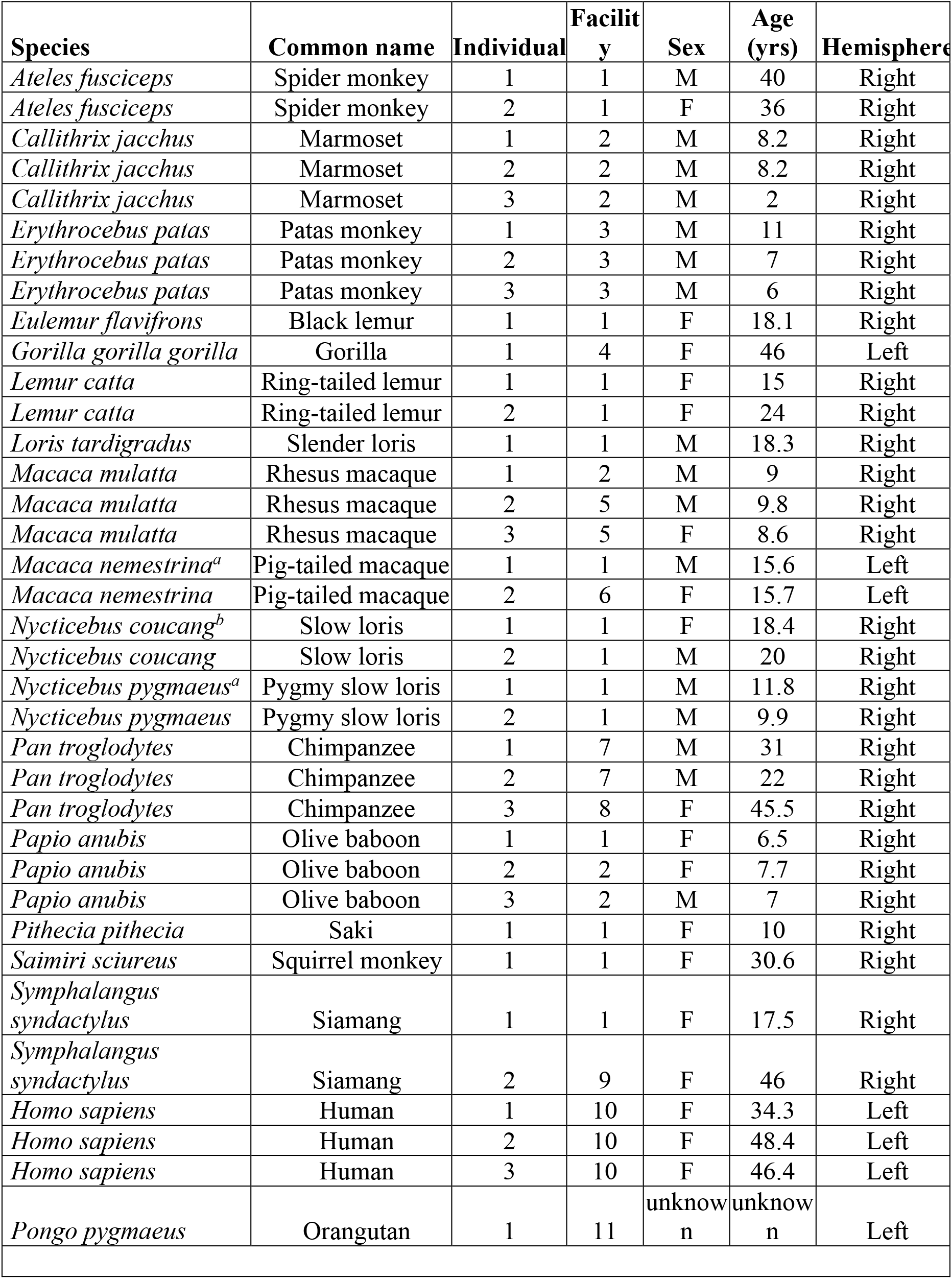
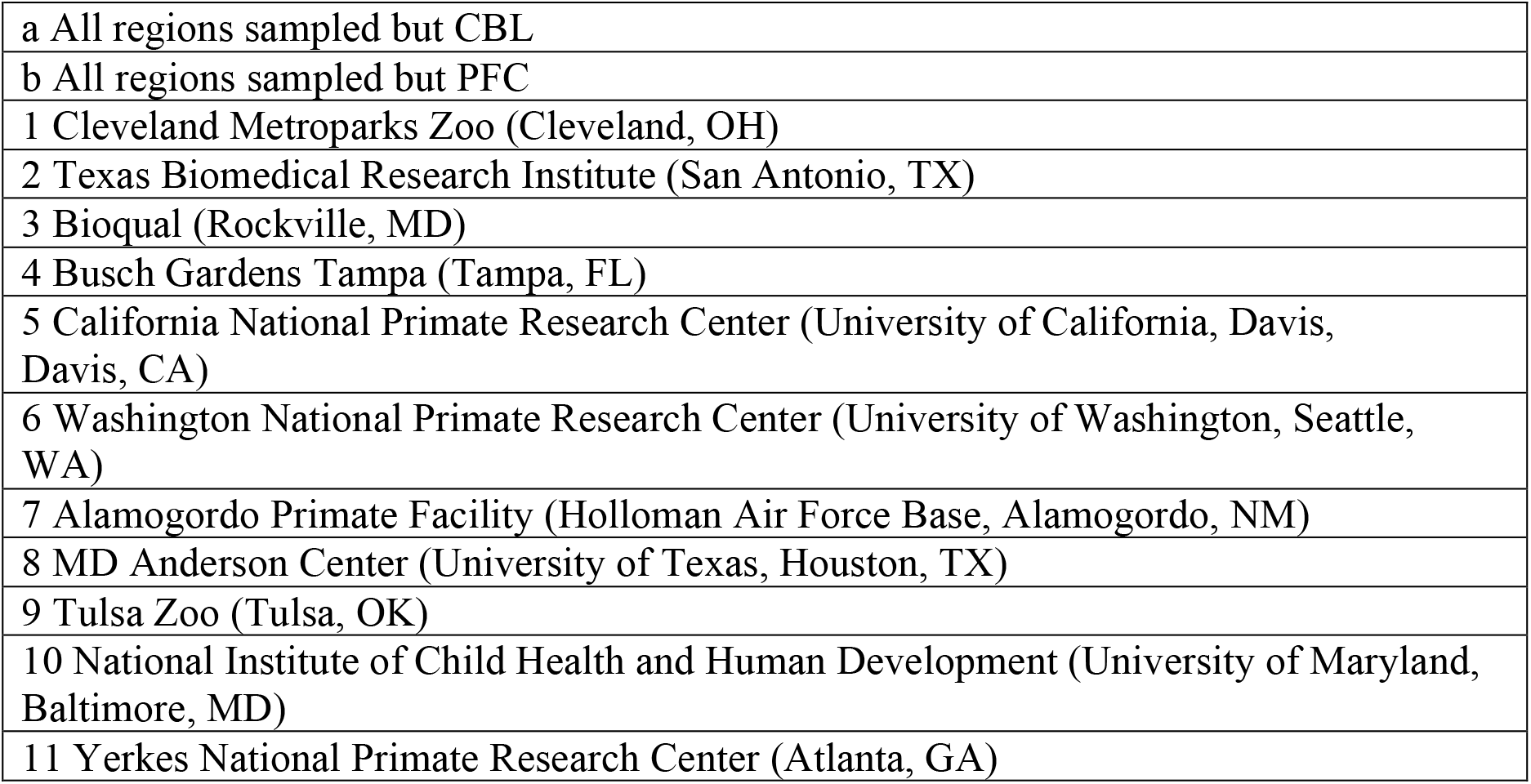
Species and sample information.

| Species | Common name | Individual | Facility | Sex | Age (yrs) | Hemisphere |
| --- | --- | --- | --- | --- | --- | --- |
| *Ateles fusciceps* | Spider monkey | 1 | 1 | M | 40 | Right |
| *Ateles fusciceps* | Spider monkey | 2 | 1 | F | 36 | Right |
| *Callithrix jacchus* | Marmoset | 1 | 2 | M | 8.2 | Right |
| *Callithrix jacchus* | Marmoset | 2 | 2 | M | 8.2 | Right |
| *Callithrix jacchus* | Marmoset | 3 | 2 | M | 2 | Right |
| *Erythrocebus patas* | Patas monkey | 1 | 3 | M | 11 | Right |
| *Erythrocebus patas* | Patas monkey | 2 | 3 | M | 7 | Right |
| *Erythrocebus patas* | Patas monkey | 3 | 3 | M | 6 | Right |
| *Eulemur flavifrons* | Black lemur | 1 | 1 | F | 18.1 | Right |
| *Gorilla gorilla gorilla* | Gorilla | 1 | 4 | F | 46 | Left |
| *Lemur catta* | Ring-tailed lemur | 1 | 1 | F | 15 | Right |
| *Lemur catta* | Ring-tailed lemur | 2 | 1 | F | 24 | Right |
| *Loris tardigradus* | Slender loris | 1 | 1 | M | 18.3 | Right |
| *Macaca mulatta* | Rhesus macaque | 1 | 2 | M | 9 | Right |
| *Macaca mulatta* | Rhesus macaque | 2 | 5 | M | 9.8 | Right |
| *Macaca mulatta* | Rhesus macaque | 3 | 5 | F | 8.6 | Right |
| *Macaca nemestrina*<sup>a</sup> | Pig-tailed macaque | 1 | 1 | M | 15.6 | Left |
| *Macaca nemestrina* | Pig-tailed macaque | 2 | 6 | F | 15.7 | Left |
| *Nycticebus coucang*<sup>b</sup> | Slow loris | 1 | 1 | F | 18.4 | Right |
| *Nycticebus coucang* | Slow loris | 2 | 1 | M | 20 | Right |
| *Nycticebus pygmaeus*<sup>a</sup> | Pygmy slow loris | 1 | 1 | M | 11.8 | Right |
| *Nycticebus pygmaeus* | Pygmy slow loris | 2 | 1 | M | 9.9 | Right |
| *Pan troglodytes* | Chimpanzee | 1 | 7 | M | 31 | Right |
| *Pan troglodytes* | Chimpanzee | 2 | 7 | M | 22 | Right |
| *Pan troglodytes* | Chimpanzee | 3 | 8 | F | 45.5 | Right |
| *Papio anubis* | Olive baboon | 1 | 1 | F | 6.5 | Right |
| *Papio anubis* | Olive baboon | 2 | 2 | F | 7.7 | Right |
| *Papio anubis* | Olive baboon | 3 | 2 | M | 7 | Right |
| *Pithecia pithecia* | Saki | 1 | 1 | F | 10 | Right |
| *Saimiri sciureus* | Squirrel monkey | 1 | 1 | F | 30.6 | Right |
| *Symphalangus syndactylus* | Siamang | 1 | 1 | F | 17.5 | Right |
| *Symphalangus syndactylus* | Siamang | 2 | 9 | F | 46 | Right |
| *Homo sapiens* | Human | 1 | 10 | F | 34.3 | Left |
| *Homo sapiens* | Human | 2 | 10 | F | 48.4 | Left |
| *Homo sapiens* | Human | 3 | 10 | F | 46.4 | Left |
| *Pongo pygmaeus* | Orangutan | 1 | 11 | unknown | unknown | Left |

|  |
| --- |
| a All regions sampled but CBL |
| b All regions sampled but PFC |
| 1 Cleveland Metroparks Zoo (Cleveland, OH) |
| 2 Texas Biomedical Research Institute (San Antonio, TX) |
| 3 Bioqual (Rockville, MD) |
| 4 Busch Gardens Tampa (Tampa, FL) |
| 5 California National Primate Research Center (University of California, Davis, Davis, CA) |
| 6 Washington National Primate Research Center (University of Washington, Seattle, WA) |
| 7 Alamogordo Primate Facility (Holloman Air Force Base, Alamogordo, NM) |
| 8 MD Anderson Center (University of Texas, Houston, TX) |
| 9 Tulsa Zoo (Tulsa, OK) |
| 10 National Institute of Child Health and Human Development (University of Maryland, Baltimore, MD) |
| 11 Yerkes National Primate Research Center (Atlanta, GA) |

### Library preparation and sequencing

RNA-Seq libraries were made using the NEBNext® mRNA Library Prep Reagent Set for Illumina. Libraries were prepared in batches of 4-8 samples of randomly sampled species and brain regions. Library sizes were checked on the Bioanalyzer (Agilent). RNA-Seq libraries were multiplexed on the NextSeq500 (Illumina) in the Genomics Resource Laboratory at the University of Massachusetts Amherst, also randomly distributed across NetSeq500 runs. All fastq files have been submitted to the SRA: https://dataview.ncbi.nlm.nih.gov/object/PRJNA639850?reviewer=136ah9q3o5ok7a7gq8jd4hhdfc

**Table S2.**
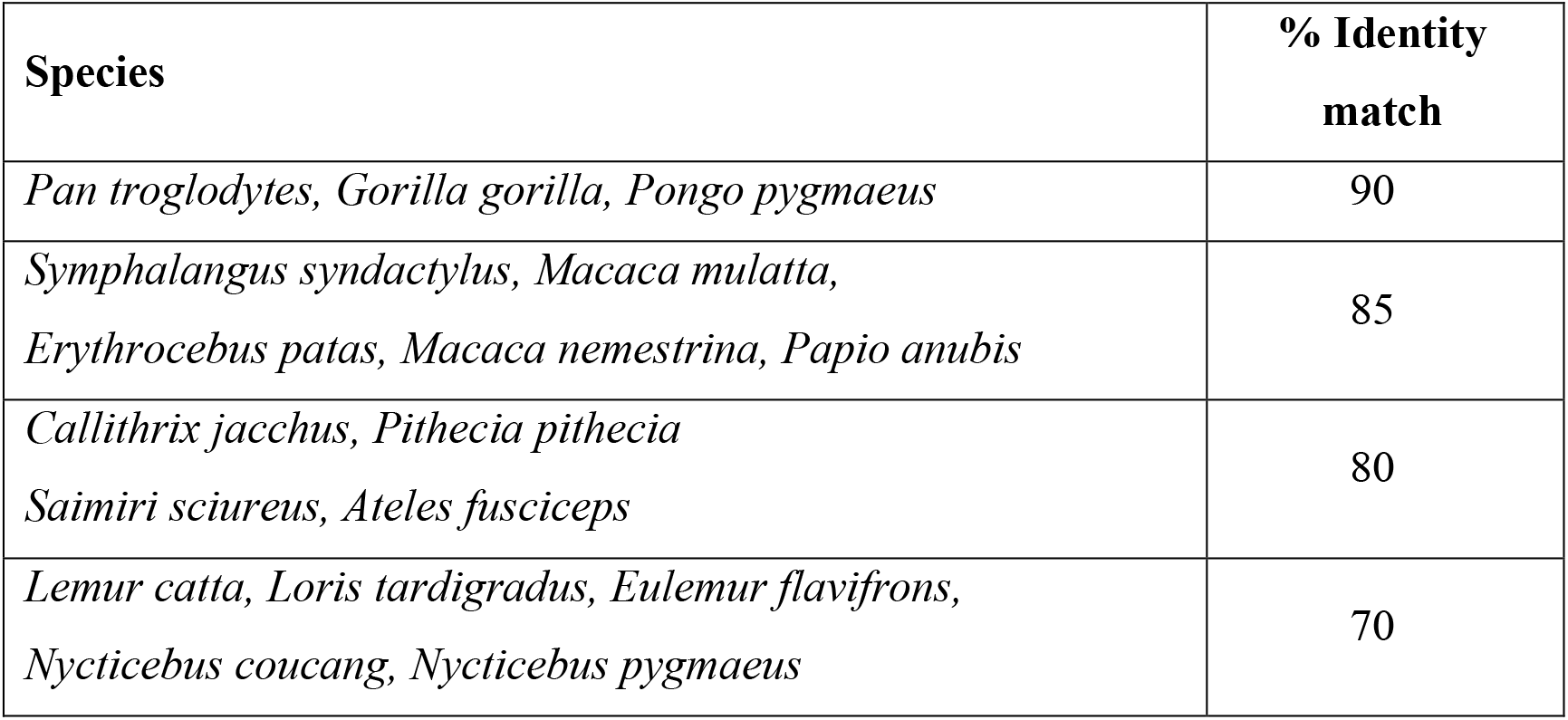
Percent identity thresholds used for orthology assignments for each species.

### Mapping and Differential Expression Analyses

Sequencing reads were assembled into species-specific transcriptomes (containing the reads from all four regions) using Trinity ^42^. These transcripts were then blasted against the human blast nt database, with the alignment thresholds for the top hits from different clades listed in **Table S2**, similar to approach in Perry, et al. ^32^. Individual libraries were then mapped to the species-specific transcriptome using bowtie ^46^ in RSEM, and counts tables were generated using RSEM ^43^. Orthology assignments were additionally checked using the Ensemble orthology alignments as a guide for the subset of species with a publicly available genome.

Counts were filtered and normalized using edgeR ^47^, with any multispecies comparisons using the GLM functionality ^44^. Gene Ontology enrichments were performed using the DAVID gene ontology tool 6.8 ^48,49^ and g:Profiler ^50^. **Table S3** shows results from ordered g:profiler enrichments (g:GOSt) performed on DE genes where q < 0.05 (note, this is not ranked on polarity of expression, just absolute change).

### Distance-based data analyses (PCoA and phenograms)

We performed a principle coordinates analyses (PCoA) based on a pairwise distance matrix of all 137 samples. The distance matrix was comprised of the top 500 most variably expressed protein-coding genes by standard deviation across samples. Pairwise distances were calculated by log2 fold change, providing a symmetrical representation of the expression ratio centered around 0 (ie. log2(2) = 1 while log2(0.5) = −1). Creating the distance matrix and plotting the PCoA were performed using the plotMDS function in the edgeR package in R. Although variation is represented across more than 20 axes (**Table S3**), the first three axes were plotted to compare patterns across primate taxa and brain region sampled (**Figure 2**). Polygons overlap the data points representing taxa or brain region. The area of each polygon was computed using functions in the sp package of R. The chull() function was used to define the points around the perimeter of each polygon, and the Polygon() function calculated the area of each. Relative areas of each polygon are listed in **Table S4. Extended Data Figure 3** displays the same data as the PCoA but uses an array of colors allowing the data from each individual species to be visualized.

**Table S3.** Eigenvalues, percent variance, and cumulative variance across disparate axes of the PCoA of the 500 most variable genes from all data sampled.

| PCoA Axis | Eigenvalue | % variance | Cumulative variance |
| --- | --- | --- | --- |
| 1 | 86.30 | 12.82 | 12.82 |
| 2 | 69.36 | 10.30 | 23.12 |
| 3 | 63.08 | 9.37 | 32.48 |
| 4 | 48.66 | 7.23 | 39.71 |
| 5 | 43.38 | 6.44 | 46.15 |
| 6 | 35.49 | 5.27 | 51.42 |
| 7 | 34.14 | 5.07 | 56.49 |
| 8 | 29.36 | 4.36 | 60.85 |
| 9 | 25.78 | 3.83 | 64.68 |
| 10 | 24.87 | 3.69 | 68.38 |
| 11 | 21.45 | 3.19 | 71.56 |
| 12 | 19.84 | 2.95 | 74.51 |
| 13 | 18.27 | 2.71 | 77.22 |
| 14 | 16.98 | 2.52 | 79.74 |
| 15 | 16.08 | 2.39 | 82.13 |
| 16 | 11.57 | 1.72 | 83.85 |
| 17 | 9.67 | 1.44 | 85.29 |
| 18 | 8.69 | 1.29 | 86.58 |
| 19 | 7.91 | 1.17 | 87.75 |
| 20 | 7.46 | 1.11 | 88.86 |
| 21 | 6.38 | 0.95 | 89.80 |
| 22 | 5.00 | 0.74 | 90.55 |

**Table S4.** Area occupied by each polygon representing either taxa or brain region in the PCoA of the 500 most variable genes from all data sampled.

|  |  | **Relative area**<br>**of multivariate**<br>**space occupied**<br>**by taxa** |  |  | **Relative area**<br>**of multivariate**<br>**space occupied**<br>**by brain**<br>**region** |
| --- | --- | --- | --- | --- | --- |
| **Axes 1 & 2** | Hominoids | 2.29 |  | CBL | 3.93 |
|  | Cercopithecoids | 1.07 |  | HIP | 4.27 |
|  | Platyrrhines | 1.18 |  | PFC | 4.65 |
|  | Strepsirrhines | 1.85 |  | V1 | 5.26 |
| **Axes 1 & 3** | Hominoids | 3.62 |  | CBL | 2.32 |
|  | Cercopithecoids | 1.56 |  | HIP | 2.32 |
|  | Platyrrhines | 1.69 |  | PFC | 2.55 |
|  | Strepsirrhines | 1.85 |  | V1 | 3.56 |
| **Axes 2 & 3** | Hominoids | 1.36 |  | CBL | 1.38 |
|  | Cercopithecoids | 0.74 |  | HIP | 1.44 |
|  | Platyrrhines | 1.69 |  | PFC | 1.55 |
|  | Strepsirrhines | 2.69 |  | V1 | 3.16 |

The same log2 fold change distance matrix was then used to create phenograms representing the similarity of gene expression profiles among samples. The neighbor-joining function in the ‘ape’ package of R created a tree based on the method proposed by Saitou and Nei (1987). The boot.phylo function estimated the reliability of given nodes of the tree by resampling over 1000 iterations. Although our objective in this analysis was to investigate patterns of evolution across the primate order, our sample included multiple individuals from the same species. By treating these samples separately, our analyses represent both within- and between-species variability.

### Phylogenetic Tree and Evolutionary Distance Analysis

Categorical enrichments for the contrasts between species and clades we are in **Table S5**. The phylogenetic tree for primates was downloaded from the UCSC Genome Browser (30 primate species)^51^. Distances between species were extracted using the Environment for Tree Exploration Toolkit^52^. The residuals and mean squared expression differences of all orthologs across 18 species were found using the package EVEE^33^, and in all contrasts, humans were used as the reference species. We then analyzed the subset of genes showing either broadly defined conserved (low) or neutral (higher) variation across species (low < q = 0.05, high q > 0.05), with categorical enrichments for these two groups in **Table S6**.

### Comparisons of the heterogenous tissues used and single cell gene expression data

In any study that derives results from homogenized tissue samples, the composition heterogeneity of the samples may drive differences in gene expression ^53^. To address this issue, we compared the expression of our tissue samples to recent studies that have performed single cell RNA-seq on neurons and astrocytes. RNA-Seq data from primary neurons and astrocytes were obtained from NCBI’s Gene Expression Omnibus (GEO) and processed in the same manner as the tissue-samples for all human samples. These included four hippocampal astrocytes, four cortical astrocytes, and one cortical neuron from Zhang et al., 2016^54^ (GEO accession number GSE73721) and three pyramidal neuron samples isolated from an unspecified brain region by the ENCODE project^55,56^ (accession numbers GSM2071331, GSM2071332 and GSM2071418). Only genes with counts greater than zero in all samples and (CPM) > 1 in all 23 samples were included in this analysis (n=7,111). A PCoA was made from a distance matrix of the top 500 most variably expressed genes by pairwise biological coefficient of variation across samples. Creating the distance matrix and plotting the PCoA were performed using the plotMDS function in the edgeR package in R. The PCoA of our human samples in comparison to primary neurons and astrocytes suggests that our heterogeneous tissue samples are not biased to contain more neurons or astrocytes as compared to each other (i.e. one tissue is not biased within this small sample set)(**Extended Data Figure 5**), and is consistent with other neural cell and brain tissue comparisons ^57^.

### Brain size analyses

Average species endocranial volumes (ECVs) were obtained primarily from Kamilar and Cooper ^58^ and Isler et al.^59^ from the mean of male and female volumes. ECVs were used since reliable brain size data does not exist for all species sample. The data for human ECV (also averaged from male and female data points) was previously published by Coqueugniot and Hublin ^60^. Isler et al. ^59^ reported minimal error when ECV was transformed to brain mass using the correction factor of 1.036g/ml ^61^, and we used this conversion to obtain brain mass estimates from ECVs for all species (**Table S7**).

**Table S7.**
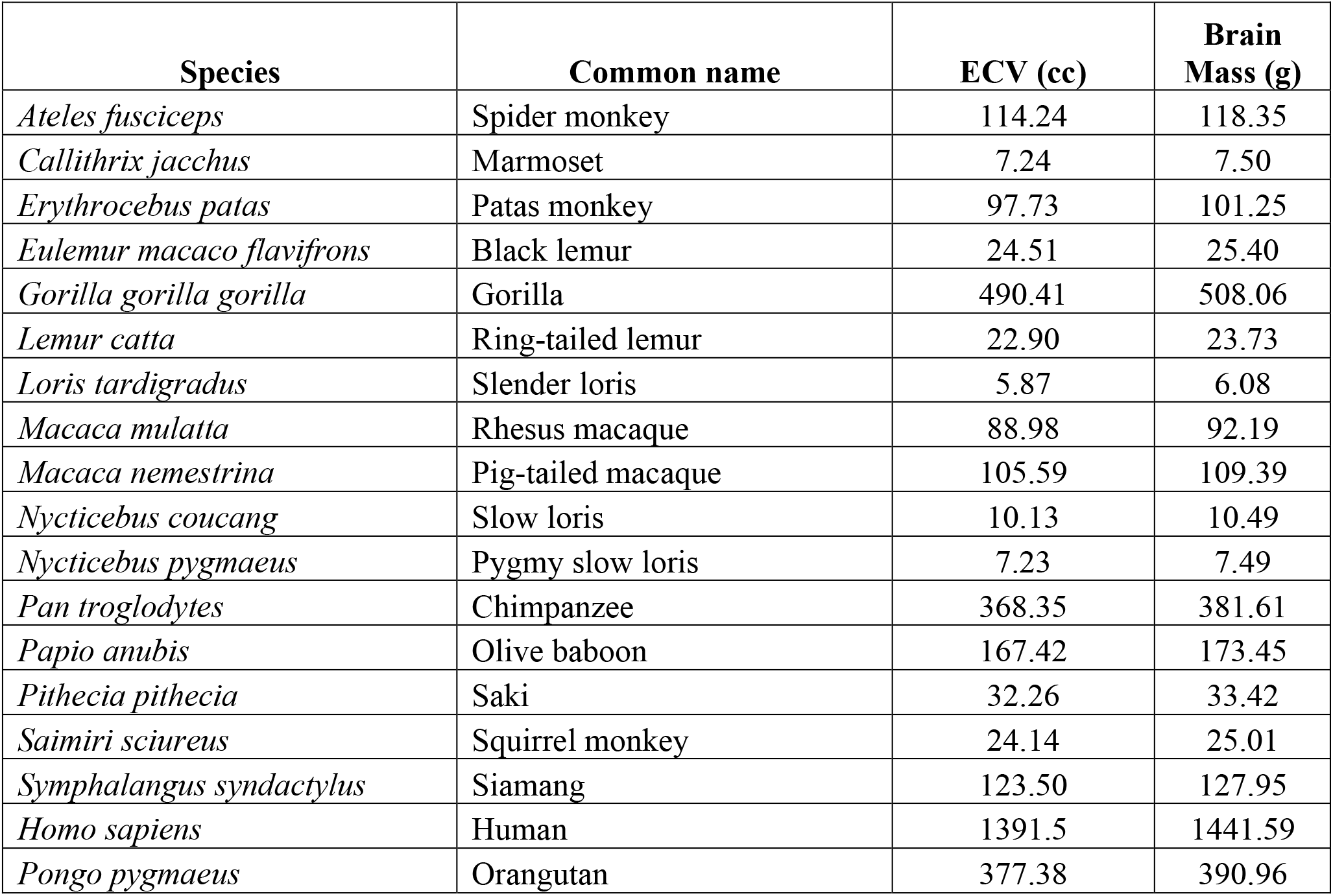
Species endocranial volumes (ECVs) were obtained from Isler et al.^59^ and Coqueugniot and Hublin^60^. Reported volumes are averages of male and female data. ECVs are transformed to brain mass using the Stephan^61^ correction factor of 1.036 g/ml.

| Species | Common name | ECV (cc) | Brain Mass (g) |
| --- | --- | --- | --- |
| *Ateles fusciceps* | Spider monkey | 114.24 | 118.35 |
| *Callithrix jacchus* | Marmoset | 7.24 | 7.50 |
| *Erythrocebus patas* | Patas monkey | 97.73 | 101.25 |
| *Eulemur macaco flavifrons* | Black lemur | 24.51 | 25.40 |
| *Gorilla gorilla gorilla* | Gorilla | 490.41 | 508.06 |
| *Lemur catta* | Ring-tailed lemur | 22.90 | 23.73 |
| *Loris tardigradus* | Slender loris | 5.87 | 6.08 |
| *Macaca mulatta* | Rhesus macaque | 88.98 | 92.19 |
| *Macaca nemestrina* | Pig-tailed macaque | 105.59 | 109.39 |
| *Nycticebus coucang* | Slow loris | 10.13 | 10.49 |
| *Nycticebus pygmaeus* | Pygmy slow loris | 7.23 | 7.49 |
| *Pan troglodytes* | Chimpanzee | 368.35 | 381.61 |
| *Papio anubis* | Olive baboon | 167.42 | 173.45 |
| *Pithecia pithecia* | Saki | 32.26 | 33.42 |
| *Saimiri sciureus* | Squirrel monkey | 24.14 | 25.01 |
| *Symphalangus syndactylus* | Siamang | 123.50 | 127.95 |
| *Homo sapiens* | Human | 1391.5 | 1441.59 |
| *Pongo pygmaeus* | Orangutan | 377.38 | 390.96 |

Within each brain region, we performed a principal components analysis (PCA) on the species average gene expression of the 500 most variable genes by standard deviation, using the prcomp function in R. For each regional PCA, 14 PCs were required to account for about 90% of the variance in gene expression. We performed multiple regression analyses to determine which of the PCs could predict brain size. Using all 14 PCs accounted for at least 95% of variation in each brain region. Akaike information criterion (both backwards and forwards methods) was applied but did little to simplify the predictive model. However, it was noted that for each brain region, PC2 was the most predictive of brain size (regional adjusted R^2^ values for PC2 against brain size were: PFC, 0.42; V1, 0.56; HIP, 0.50; CBL, 0.36). The 500 genes and their loadings on PC2 are listed in **Table S8**. Across the four sampled regions, we find a remarkable degree of uniformity in the extent to which individual genes affect brain size, but the cerebellum displays the most unique signature of the regions sampled (**Figure 5**).

### Scans for positive selection in gene regulatory regions

We tested for signs of positive selection in 5kb promoter regions of the top 200 genes that were differentially expressed between human and chimpanzee or came up as significant in the correlations with brain size. Coordinates of human promoters were obtained from Gencode ^62^ or RefSeq ^63^ annotations of human genome assembly GRch38 using the 5’-most exon coordinate as the transcription start site. Genes were excluded if annotations were not available. Portions of promoters overlapping genes were removed and only the 3’ most region was used for analysis. Promoters less than 500 bp were excluded from the analysis. We used 100 kb non-coding regions centered on the promoter as a proxy for a neutral rate of substitution. Within these 100kb windows, the first introns of genes were removed since these often contain regulatory elements. Coordinates of promoters and introns were used to download human-chimpanzee-macaque alignments from the Ensembl 6 primates EPO alignment generated in Ensembl release 80 in the Compara API. Promoters were discarded if alignment data were not available for all 3 species in either promoter or neutral regions. The test for selection was performed using modified code from Haygood et al. ^35^, and the analysis pipeline is available on GitHub (https://github.com/jpizzollo/Selection-Workflow), and run using HyPhy software ^45^. P-values were used to identify promoters with significantly higher rates of substitution on the human branch (**Table S9**). Promoters were also tested for selection using phyloP ^64^ to look for signs of selection on the human branch of the human, chimpanzeee, and macaque tree generated from the Ensembl 6 primate EPO alignment. P-values for acceleration were generated for full promoter sequences using the SPH method, and for individual nucleotides using the LRT method ^64^ (**Table S10**).

## List of all Supplemental Tables

**Table S1**. Species and sample information.

**Table S2**. Percent identity thresholds used for orthology assignments for each species.

**Table S3**. Eigenvalues, percent variance, and cumulative variance across disparate axes of the PCoA of the 500 most variable genes from all data sampled.

**Table S4**. Area occupied by each polygon in the PCoA of the 500 most variable genes from all data sampled.

**Table S5**. Gene ontology categorical enrichments (Molecular Function (MF), Biological Process (BP), and Cellular Component (CC)). Species comparisons included are 1. Human-Chimpanzee, Human-Siamang, Human-Rhesus Macaque, Human-Marmoset, and Human-Lemur. Clade comparisons are for Haplorrhines vs Strepsirrhines.

**Table S6**. GO Biological Process categorical enrichments (p < 0.05) from the evolutionary rate tests.

**Table S7**. Species endocranial volumes (ECVs) were obtained from Isler et al. (2008) and Coqueugniot and Hublin (2012). Reported volumes are averages of male and female data. ECVs are transformed to brain mass using the Stephan (1960) correction factor of 1.036 g/cc.

**Table S8**. Genes and their loadings for PC2, the axis which is most strongly correlated with brain size in each brain region.

**Table S9**. Full table from the scans for selection. Results from test for positive selection using the Haygood et al. method in 0.5-5kb proximal regulatory regions of the top 200 DE genes between human and chimpanzee or genes correlated with differences in brain size.

**Table S10**. Expanded results from Table 1. Here the table differs from the main table in the addition of the nucleotides under selection columns generated with phyloP’s LRT method to test accelerated substitution base-by-base within an alignment.

## Notes

### Competing Interest Statement

The authors have declared no competing interest.

